# Higher-order visual areas broaden stimulus responsiveness in mouse primary visual cortex

**DOI:** 10.1101/2021.02.16.431393

**Authors:** Matthijs N. oude Lohuis, Alexis Cerván Cantón, Cyriel M. A. Pennartz, Umberto Olcese

**Author notes:** **Author List Footnotes:** Lead Contact. **Contact Info:** Correspondence.

## Abstract

Over the past few years, the various areas that surround the primary visual cortex in the mouse have been associated with many functions, ranging from higher-order visual processing to decision making. Recently, some studies have shown that higher-order visual areas influence the activity of the primary visual cortex, refining its processing capabilities. Here we studied how *in vivo* optogenetic inactivation of two higher-order visual areas with different functional properties affects responses evoked by moving bars in the primary visual cortex. In contrast with the prevailing view, our results demonstrate that distinct higher-order visual areas similarly modulate early visual processing. In particular, these areas broaden stimulus responsiveness in the primary visual cortex, by amplifying sensory-evoked responses for stimuli not moving along the orientation preferred by individual neurons. Thus, feedback from higher-order visual areas amplifies V1 responses to non-preferred stimuli, which may aid their detection.

## Introduction

Over the past few years the various areas which make up the mouse visual cortical system have emerged as a prime model to study the functional architecture underlying vision in mammals (Andermann et al., 2011; Glickfeld and Olsen, 2017; Glickfeld et al., 2014; Marshel et al., 2011; Wang and Burkhalter, 2007). The anterior and lateral borders of primary visual cortex V1 are surrounded by an array of areas, collectively called higher-order visual areas (HVAs), each having a unique connectivity pattern and visual response properties (Andermann et al., 2011; Marshel et al., 2011; Wang et al., 2012). Several studies have investigated what functions each of these areas might fulfill in visual processing. A wide range of functions has been found, complementing V1 in orientation discrimination and contrast detection (Jin and Glickfeld, 2020), spatial integration (Murgas et al., 2020), perception of higher-order visual features (Khastkhodaei et al., 2016) and illusory contours (Pak et al., 2020), object and shape representations (Matteucci et al., 2019; Tafazoli et al., 2017). Furthermore, HVAs partially overlap with the rodent posterior parietal cortex, and have been implicated in several functions beyond simple visual processing, for instance multisensory integration (Meijer et al., 2020; Olcese et al., 2013; Song et al., 2017), (multi)sensory evidence accumulation and decision making (Erlich et al., 2015; Hanks et al., 2015; Licata et al., 2017; Raposo et al., 2014) and navigation (Harvey et al., 2012; Krumin et al., 2018). Moreover, HVAs play a significant role in sensory processing by means of the input they provide not only to each other, but also to V1 (Wang et al., 2012).

Feedback projections from HVAs to V1 have been found to be functionally organized (Kim et al., 2018; Marques et al., 2018), similarly to local connections (Cossell et al., 2015; Ko et al., 2011) and feedforward projections from V1 to HVAs (Berezovskii et al., 2011; Glickfeld et al., 2013). These feedback projections have been associated with a variety of essential forms of visual processing: response facilitation (Nurminen et al., 2018; Pafundo et al., 2016), surround suppression (Nassi et al., 2013; Nurminen et al., 2018; Vangeneugden et al., 2019) and predictive processing (Keller et al., 2020). A recent study, in particular, showed that each HVA differently impacts the activity of V1 neurons based on their visual response properties (Huh et al., 2018). Inactivating either the anterolateral (AL) or posteromedial (PM) area primarily reduced responses of those V1 neurons showing functional properties similar to those of AL and PM, respectively. This study focused on tuning of V1 cells to spatial frequency and investigated how inactivating AL and PM modulates firing rate responses to drifting gratings moving along the preferred orientation of single neurons. Overall, previous studies thus indicate that feedback projections from HVAs to V1 may provide a mechanism to enhance processing of specific visual stimuli, based on the response properties of each HVA. In this study, we combined optogenetics and ensemble recordings to investigate how HVAs contribute to a broader spectrum of V1 functions beyond processing of only optimal (preferred) stimuli, such as orientation and direction selectivity, receptive field size and single-trial encoding of visual features. Surprisingly, we found that, in addition to the previously reported, functionally specific feedback (in which modulation of V1 varies based on the functional tuning of each HVA), AL and PM similarly enhance V1 responsiveness to visual stimuli, but globally diminish receptive field size and selectivity to stimulus orientation and direction. Thus, in addition to previously discovered functions, HVAs may also implement an amplification in their feedback to V1 which may aid stimulus detection.

## Results

To investigate how HVAs influence V1 responses, we focused on two areas with the largest known differences in tuning to spatial and temporal frequencies of visual stimuli: AL and PM. While AL neurons preferentially respond to visual stimuli with high temporal frequencies and low spatial frequencies, the opposite is true for area PM (Andermann et al., 2011; Marshel et al., 2011). We performed dual-area silicon probe recordings in head fixed mice from either V1 and AL or V1 and PM (Fig. 1A). Recordings were done in both the awake and anesthetized state. As recently reported (Keller et al., 2020), feedback modulation from HVAs to V1 is reduced under anesthesia, but the effect of brain state on V1 response properties is poorly understood (Olcese et al., 2018), although previous studies reported a reduction in direction tuning in isoflurane anesthesia compared to wakefulness (Goltstein et al., 2015). Localized nano-injections of a viral vector mediating Cre-dependent expression of channelrhodopsin were performed in either AL or PM of PV-Cre mice (Madisen et al., 2012) – Fig. 1A,C. Areas V1, AL and PM were localized via intrinsic optical signal imaging (IOI, Fig. 1B). Blue-light illumination was used to inactivate either area AL or PM, via over-activation of parvalbumin-positive (PV+) interneurons (Olcese et al., 2013) – Fig. 1D-G. We experimentally verified that optogenetic inactivation was confined to areas AL and PM, and did not affect V1 directly (Fig. 1C,G). Specifically, inactivation of either AL or PM greatly reduced the activity of putative excitatory neurons in the illuminated area (Fig. 1G, top), but only had a minor effect of spontaneous firing activity in V1 (Fig. 1G, bottom). Moving bars were used as visual stimuli to evoke activity in V1 and HVAs (Fig. 1H). Compared to drifting gratings, moving bars enable to assess receptive field sizes of the recorded neurons (Niell and Stryker, 2008). Bars were moving over 8 different orientations at three different speeds, namely those that previous studies indicated as being preferred by PM (20 deg/s), V1 (40 deg/s) and AL (70 deg/s) (Andermann et al., 2011; Marshel et al., 2011). To verify this speed preference, we computed, separately for each neuron and area, the response to a bar moving along the preferred orientation, independently for each speed. For each neuron, responses were normalized to the speed evoking the strongest response (Fig. 1I). For recordings performed under anesthesia, the speed preference of neurons in each area was in line with the literature (Fig. 1I-right). During wakefulness we found a shift for all areas to lower preferred speeds compared to anesthesia (Fig. 1I-left), although the differential speed preference of neurons in area PM and AL – with area PM selective for low speeds and AL for faster speeds – was preserved. Optogenetic inactivation of either area PM or AL suppressed sensory-evoked responses in the illuminated area – PM or AL, respectively (Fig. S1A,B). Sensory-evoked responses in V1 were not suppressed, but only reduced, by inactivation of one HVA (Fig. 1J). Such reduction was limited to sensory-evoked responses, with no effect on spontaneous activity (Fig. 1G), and can be interpreted in the first instance to be the consequence of impaired recurrent connectivity from AL or PM (see also STAR Methods).

**Figure 1.**
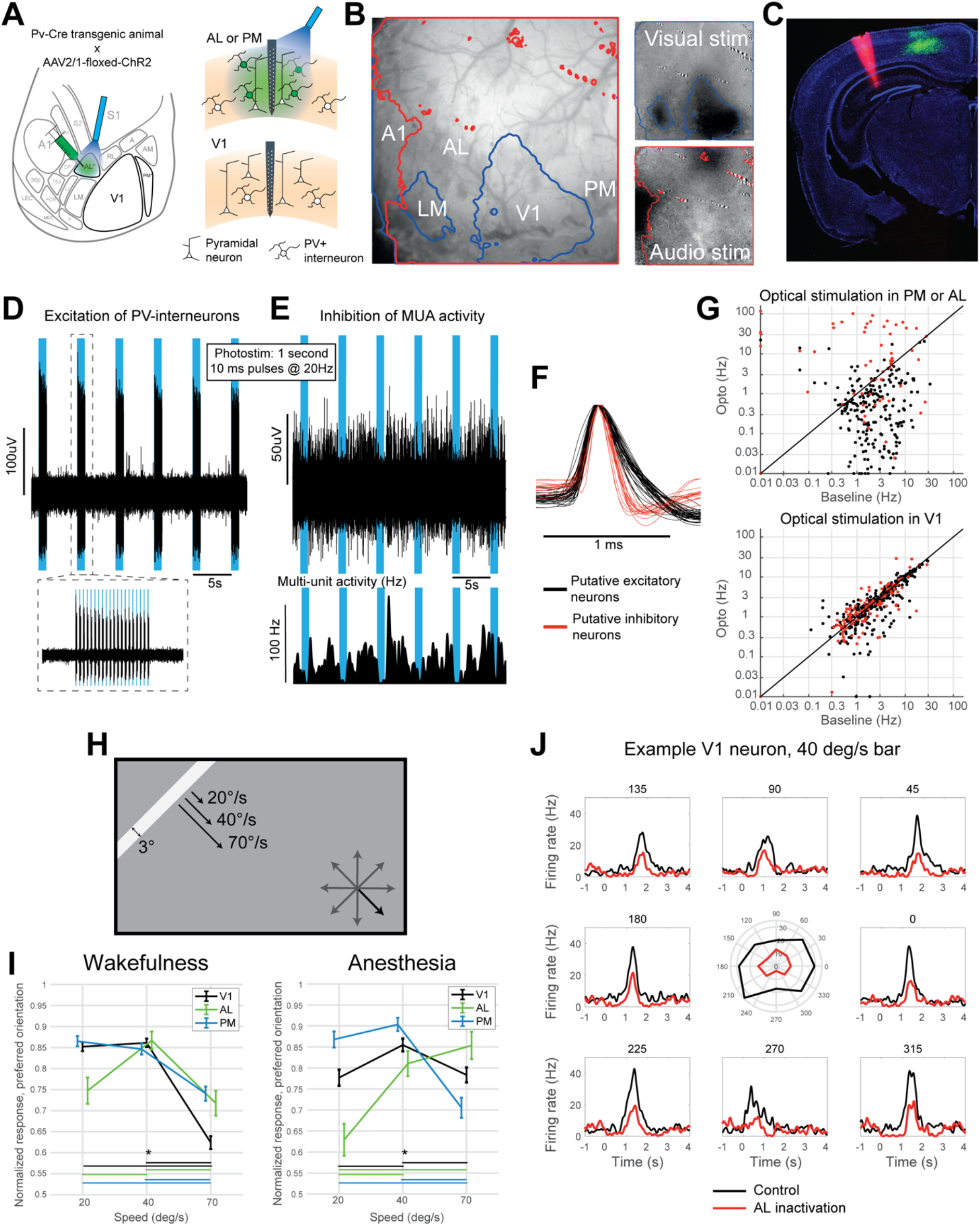
Experimental setup. **A.** Schematic of the experimental design. Left: top view of the left cortical hemisphere of a mouse, with subdivision in cortical areas - based on (Wang and Burkhalter, 2007). Adeno-associated viral vector mediating the Cre-dependent expression of ChR2 was injected in area AL (or PM, not shown). During experiments an optic fiber (blue) was placed on top of AL (or PM) to over-activate Cre-expressing PV+ interneurons and inactivate area AL (or PM). Right: scheme of coronal sections of either AL/PM (top) or V1 (bottom) showing laminar probe recordings in both areas. Expression of ChR2 and fiber-optic-mediated illumination were confined to area AL or PM (top). **B.** Intrinsic signal imaging was used to localize cortical areas. Visual stimuli (top right) and auditory stimuli (bottom right) were used to activate and thus identify the location of visual and auditory cortices. The borders of visually- and auditory-evoked signals (blue and red curves, respectively) were overlaid on the vessel map (left) to identify the location of V1, AL and PM. In this example, V1, LM and A1 were directly activated by visual or auditory stimuli. The location of AL and PM was determined based on published maps of the mouse visual system (see *A).* **C.** Coronal section showing ChR2-conjugated GFP expression in area PM (green) and the location of a laminar probe positioned in V1 and stained with DiI (red). **D.** Example neuronal trace from a PV+ interneuron recorded in area PM during optogenetic illumination in control trials (during awake recordings). Raw trace from a channel showing spiking activity evoked in a PV+ interneuron by optogenetic illumination. Blue areas indicate 1 s periods in which the blue laser was on. The inset shows the pattern of optogenetic illumination (10 ms ON - 40 ms OFF) during each illumination period. **E.** Top: Example multiunit activity (MUA) trace recorded in area PM during optogenetic illumination in control trials. In contrast with panel *D*, spiking activity decreased during illumination periods. Bottom: Firing rate traces as extracted from MUA activity shown above. Notice the decrease in firing rates during illumination. **F.** Average action potential waveforms from a selection of putative excitatory neurons (black, characterized by broad spikes) and putative inhibitory neurons (red, characterized by narrow spikes). **G.** Scatter plots of firing rates of individual neurons during spontaneous baseline activity and during optogenetic stimulation in areas PM or AL (top) and V1 (bottom), for both putative excitatory and inhibitory neurons (black and red points, respectively). Top: average spontaneous firing rates for putative excitatory neurons significantly decreased (mean values: 3.5 Hz and 1.3 Hz, p<0.001, paired t-test), while those for putative inhibitory neurons significantly increased (mean values: 4.5 Hz and 20.9 Hz, p<0.001, paired t-test) upon optogenetic stimulation of PM and AL, for neurons located in area PM and AL. Bottom: average spontaneous firing rates for putative excitatory neurons significantly decreased (mean values: 3.9 Hz and 3.5 Hz, p<0.001, paired t-test), while those for putative inhibitory neurons were not affected (mean values: 3.1 Hz and 3.2 Hz, p>0.2, paired t-test) upon optogenetic stimulation of PM and AL, for neurons located in V1. Although firing rates of putative excitatory neurons in V1 decreased upon optogenetic stimulation, such decrements were much weaker than those reported in PM and AL. This is in line with a loss of excitatory drive in V1 due to the inactivation of areas AL or PM. **H.** Outline of the visual stimuli (moving bars moving at different speed along 8 possible directions). **I.** Speed preference for neurons located in V1 (black), PM (blue) and AL (green) as a function of brain state (left: wakefulness; right: anesthesia). For each neuron, responses to the preferred orientation were computed across the three bar speeds, and normalized to the highest response (corresponding to the preferred bar speed). Asterisks indicate significant differences between speeds, for neurons located in the same area (p<0.05, one-way Anova with post-hoc Tukey test). **J.** PSTHs computed for an example neuron in V1, for bars moving at 40 deg/s across the 8 different orientations, with and without optogenetic stimulation (black and red traces, respectively). The polar plot at the center of the panel shows the tuning curve of the example neuron. See also Fig. S1.

### Inactivation of areas AL and PM globally decreases V1 responses to moving bars

Having established that areas AL and PM show sensory-evoked responses which differ based on the specific speed at which presented bars move (Fig. 1I), we wondered whether inactivation of AL and PM would differentially modulate V1 responses to bars moving at different speeds. Surprisingly, we found that silencing either AL or PM consistently reduced V1 responses to bars moving at all the speeds we tested, and for both preferred and nonpreferred orientations (Fig. 2A-D). This was the case during both awake (Fig. 2A-D) and anesthetized (Fig. S1C,D) recordings. We next asked if the extent to which V1 responses were reduced varied as a function of bar speed. For each neuron and bar speed, we computed the relative change in the response to a bar moving along the neuron’s preferred direction following optogenetic inactivation of either area AL or PM. No significant difference was found between speeds when inactivating either AL or PM (Fig. 2E). Only for bars moving at 20 deg/s, we found than PM inactivation reduced V1 responses more strongly than AL inactivation. To further explore the possible occurrence of a functionally specific effect, we subdivided V1 neurons in three groups based on the bar speed for which they showed the highest response - cf. (Huh et al., 2018) - and assessed whether AL and PM inactivation had different effects for the three groups of neurons. While we found no significant differences for bars (Fig. 2F), we did find some differences when presenting drifting gratings, in line with (Huh et al., 2018) - see Fig. S1G.

**Figure 2.**
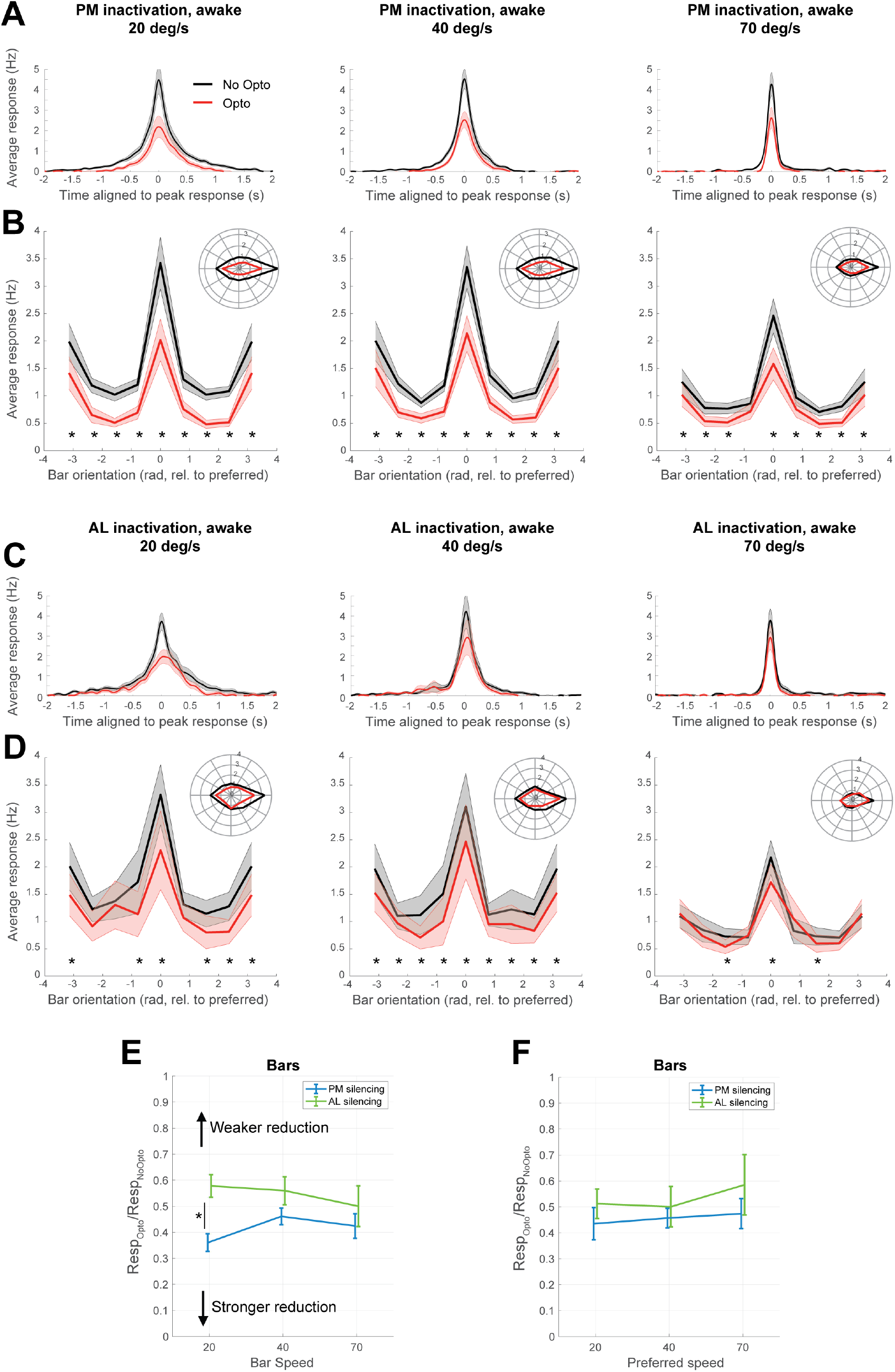
Optogenetic inactivation of area AL and PM during wakefulness depresses sensory-evoked responses in V1. **A.** PSTHs (aligned to peak latency) averaged over all V1 neurons responding to visual stimuli during awake recordings in the absence or presence of optogenetic inactivation of area PM (black and red curves, respectively). Curves with shading indicate mean ± SEM. From left to right: responses to bars moving at 20, 40 and 70 deg/s. **B.** Tuning curves of average responses of V1 neurons during awake recordings to bars moving at different orientation (aligned by the preferred orientation, which is set at 0 rad) in the absence or presence of optogenetic inactivation of area PM (black and red curves, respectively). Curves with shading indicate mean ± SEM. Inset: Average response displayed in polar coordinates. Asterisks indicate significant differences between responses to bars moving at a given orientation in the absence or presence of PM inactivation (p<0.05, paired t-test, FDR-corrected). From left to right: responses to bars moving at 20, 40 and 70 deg/s. **C.** Same as *A*, but now for the inactivation of area AL. **D.** Same as *B*, but now for the inactivation of area AL. **E.** Relative effect of optogenetic inactivation of either PM (blue) or AL (green) on V1 responses evoked by moving bars during awake recordings (Resp_Opto_/Resp_NoOpto_), as a function of bar speed. Asterisks indicate significant differences either between speeds (for a given area) or between inactivation of distinct areas, given the same speed (p<0.05, 2-way anova with post-hoc Tukey test). Error bars indicate mean ± SEM. **F.** Same as *E*, but subdividing neurons based on their preferred speed, and only considering the effect of optogenetic inactivation on the preferred speed. See also Fig. S1.

In conclusion, inactivating both AL and PM generally decreased V1 responses to moving bars. While we were able to confirm the previously reported presence of a functionally specific effect of AL and PM silencing on V1 activity (i.e., being dependent on the speed preference of each HVA), this effect was weaker than the generalized decrease in responses observed across speeds (Fig. 2E-F, Fig. S1G).

### Inactivation of AL and PM reduces receptive field size of V1 neurons

The general reduction in visually evoked responses (Fig. 1J, 2A,C) made us wonder whether inactivating areas PM and AL would also reduce the receptive field size of V1 neurons. Indeed, in awake recordings we observed a reduction of receptive field size, which was especially pronounced for bars moving at low speed (20 deg/s; Fig. 3A,B), but still present for bars moving at higher speeds, albeit only at some orientations (Fig. 3A,B). Results were very similar for recordings performed under anesthesia, with the main difference being larger receptive fields for V1 neurons in anesthetized than in awake recordings (Fig. S2). Thus, inactivating AL and PM not only reduces V1 responses to moving bars but, especially for low-speed stimuli, makes them more spatially localized.

**Figure 3.**
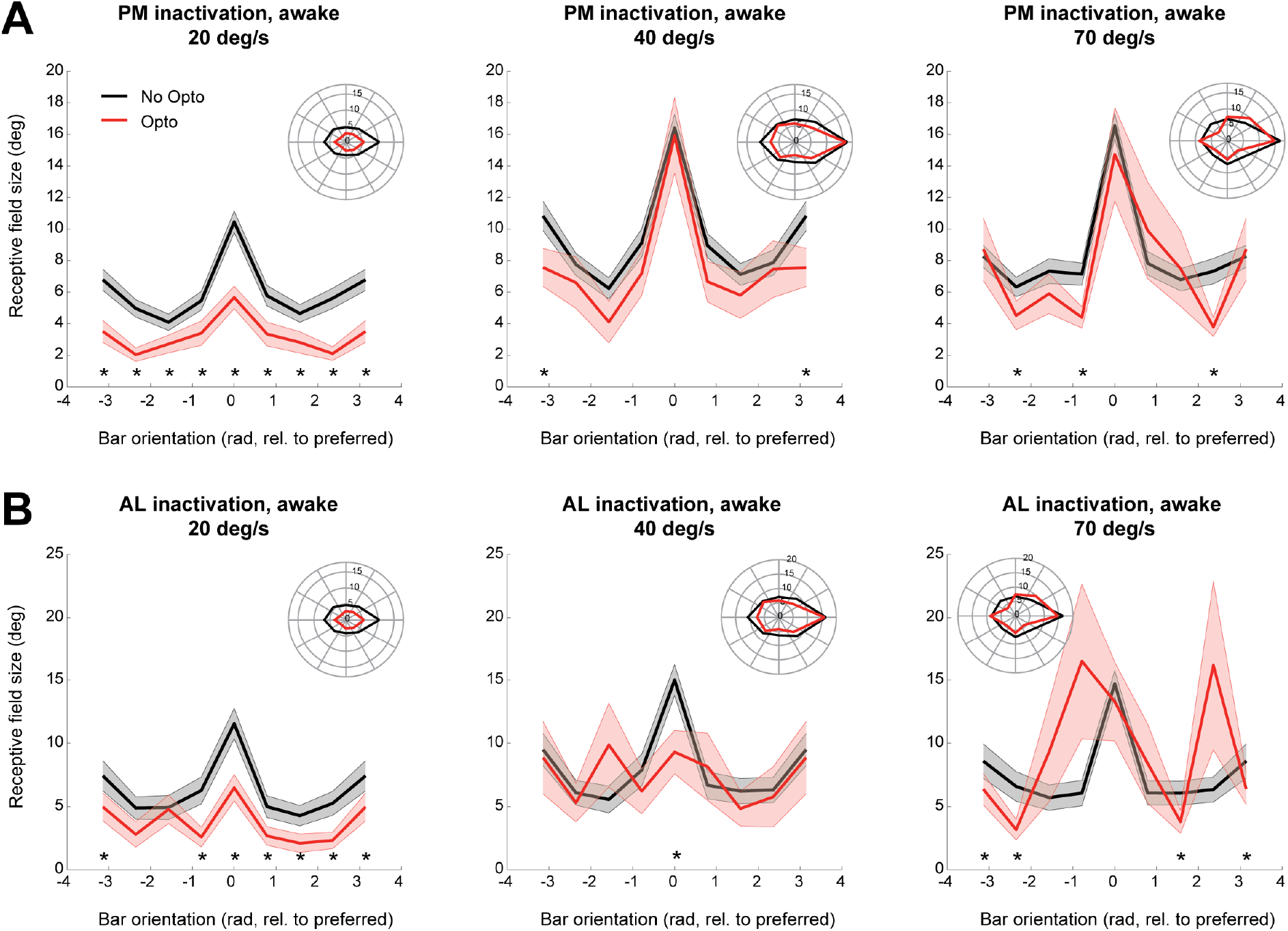
Optogenetic inactivation of area AL and PM during wakefulness reduces receptive field size of V1 neurons. **A.** Tuning curves of receptive field size of V1 neurons during awake recordings to bars moving at variable orientation (aligned by the preferred orientation, which is set at 0 rad) in the absence or presence of optogenetic inactivation of area PM (black and red curves, respectively). Curves with shading indicate mean ± SEM. Inset: average receptive field size displayed in polar coordinates; preferred orientation is aligned to the 0 deg (rightward) direction. Asterisks indicate significant differences between receptive field size to bars moving at a given orientation in the absence or presence of optogenetic inactivation of area PM (p<0.05, paired t-test, FDR-corrected). From left to right: responses to bars moving at 20, 40 and 70 deg/s. **B.** Same as *A*, but now for optogenetic inactivation of area AL. See also Fig. S2.

### Orientation and direction selectivity of V1 neurons are enhanced for moving bars when AL and PM are inactivated

To better explore the functional significance of the reduction in V1 responses and receptive field size, we wondered how silencing AL and PM might affect orientation and direction selectivity of V1 neurons. These were quantified by computing, respectively, a global orientation selectivity index (gOSI) and a global direction selectivity index (gDSI) (Ibrahim et al., 2016; Mariño et al., 2005) - see STAR Methods. Based on previous literature (Pafundo et al., 2016), we hypothesized that inactivation of HVAs would reduce both gOSI and gDSI. To our surprise, both gOSI and gDSI instead increased, across all bar speeds and in both wakefulness and anesthesia, regardless of whether AL or PM was inactivated (Fig. 4A-D, Fig. S3A-D). No significant difference was found between inactivation of AL or PM, or between bar speeds (Fig. 4E-F) - although a more prominent enhancement for both gOSI and gDSI was observed at low speeds compared to high speeds when PM was inactivated during anesthesia (Fig. S3E-F). Overall, these results suggest that inactivation of AL and PM differentially reduces responses to bars moving along preferred and non-preferred orientations, in a way that enhances orientation and direction selectivity.

**Figure 4.**
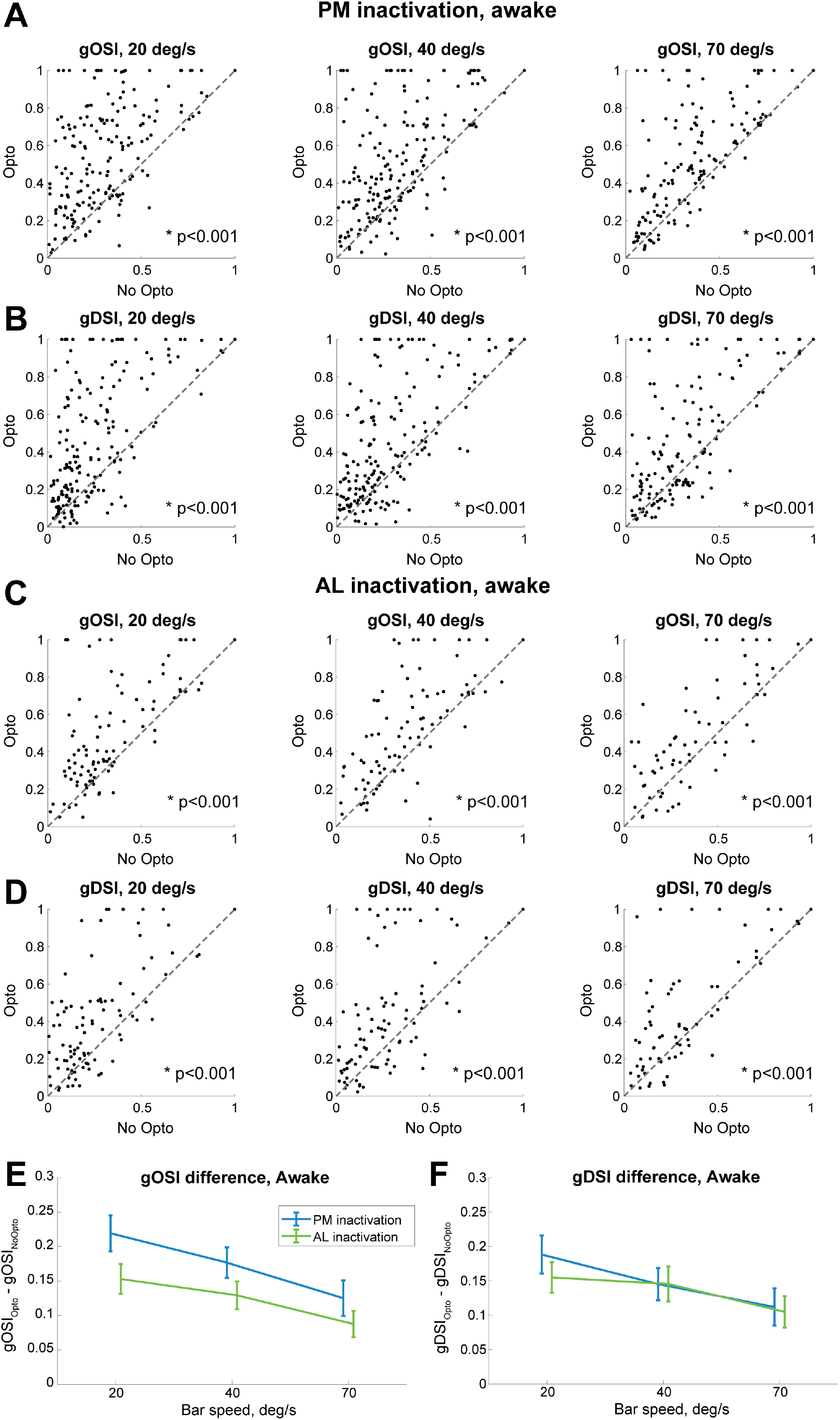
Optogenetic inactivation of area AL and PM during wakefulness enhances orientation and direction selectivity in V1. **A.** Scatter plots showing the orientation selectivity of V1 neurons during awake recordings in the absence or presence of optogenetic inactivation of area PM (x and y axis, respectively). Each point corresponds to a single neuron. Asterisks indicate significant differences between gOSI for bars moving at a given orientation in the absence or presence of optogenetic inactivation of area PM (p<0.001, paired t-test). From left to right: gOSI for bars moving at 20, 40 and 70 deg/s. **B.** Same as *A*, but now for direction selectivity (gDSI). **C.** Same as *A*, but now for optogenetic inactivation of area AL. **D.** Same as *B*, but now for optogenetic inactivation of area AL. **E.** Average change in the orientation selectivity of V1 neurons during awake recordings as a function of bar speed and area being inactivated (PM: blue; AL: green). Error bars indicate mean ± SEM. No significant difference was found, neither between bar speeds, nor between areas. **F.** Same as *E*, but now for direction selectivity. No significant difference was found, neither between bar speeds, nor between areas. See also Fig. S3.

### Single-trial decoding of the orientation of moving bars improves in V1 during wakefulness following inactivation of AL or PM

Orientation and direction selectivity indices are computed over average responses to visual stimuli. Therefore, we wondered if, in spite of enhancing gOSI and gDSI, inactivating AL and PM might have a different effect at the single-trial level. We reasoned that a reduction in response size may reduce response variability at the average level (hence the improved gOSI and gDSI), yet still impair single-trial response selectivity. To investigate this, we implemented a decoding approach to measure how well the direction of moving bars could be decoded from single-trial V1 responses (see STAR Methods). In line with the increase in gOSI and gDSI, we found that, during awake recordings, inactivating AL or PM significantly enhanced single-trial decoding of bar orientation, irrespective of bar speed, and with a stronger improvement following AL than PM inactivation (Fig. 5). Similar results were observed for recordings under anesthesia, although the improvement in decoding following inactivation of PM or AL was smaller than for awake recordings (Fig. S4). Thus, inactivation of AL and PM not only reduces visually evoked responses and makes receptive fields smaller (i.e., spatially more precise), but also - possibly by differentially affecting responses to bars moving along preferred vs. non-preferred orientations and directions - enhances the selectivity of V1 neurons to the orientation and direction of moving stimuli, both at the average and single-trial level.

**Figure 5.**
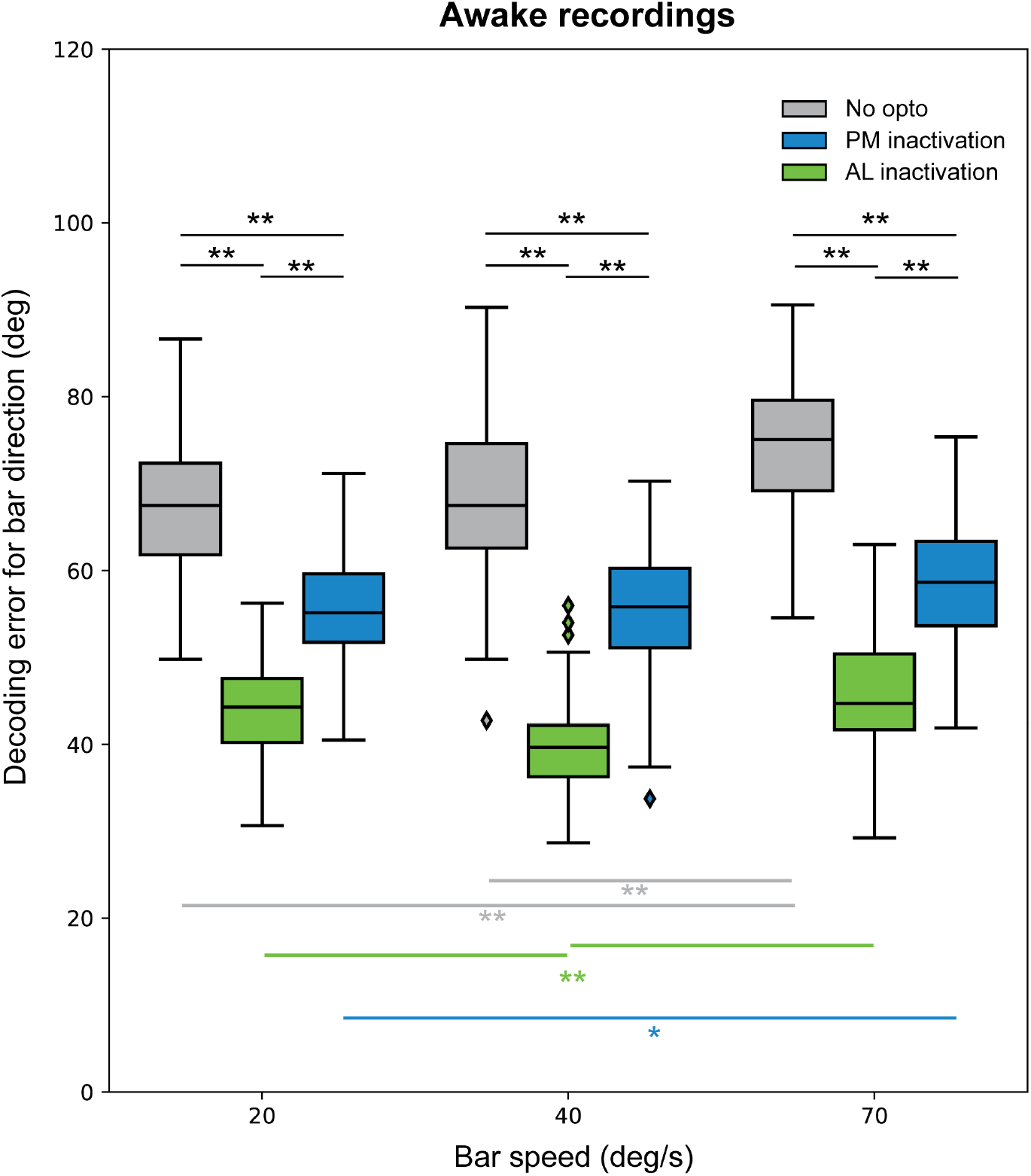
Optogenetic inactivation of area AL and PM during wakefulness enhances single-trial decoding of bar orientation in V1. Boxplots show the error (in deg) made by a decoder trained to estimate the orientation of a moving bar presented during individual trials, from the activity of a pseudo-population of V1 neurons (see STAR Methods for decoding analysis). A lower decoding error indicates better ability to decode the orientation of a moving bar. Boxplots were separately computed for trials without optogenetic stimulation (grey) and for trials in which either AL (green) or PM (blue) was inactivated. Asterisks indicate significant differences (*: p<0.05, **: p<0.001; 2-way anova with post-hoc Tukey test). See also Fig. S4.

### AL and PM provide a modulatory gain to V1, which enhances weak visual responses during wakefulness

The above results paint a counter-intuitive role of AL and PM, which would have a primarily detrimental role on the response selectivity of V1 neurons, if their sole purpose was to signal the orientation and direction of visual stimuli. We thus investigated in more depth whether inactivating AL and PM has a differential effect based on the size of a neuron’s response to a given orientation. We reasoned that the modulation provided by AL and PM onto V1 might differentially enhance responses of different sizes (such as those elicited by bars moving along preferred vs. non-preferred orientations and directions), as previously shown for other forms of cortical gain modulation (Ferguson and Cardin, 2020). If the role of AL and PM is to selectively enhance V1 responsiveness to nonpreferred orientations/directions, this might also explain the enhanced orientation and direction selectivity which result from inactivation of PM and AL. Therefore, we tested whether AL and PM might provide a form of additive or multiplicative gain modulation of V1 responses, as these are often found in visual cortex (Wilson et al., 2012). To this aim, we subtracted or divided V1 responses following optogenetic inactivation of AL and PM by V1 responses in control conditions (Fig. 6A-B). During awake recordings, AL and PM implemented neither an additive nor a multiplicative form of gain modulation. In case of additive modulation, the curves in Fig. 6A would have been flat, reflecting a similar reduction in evoked responses following inactivation of AL or PM; similarly, in case of multiplicative modulation, the curves in Fig. 6B would not have shown differences between preferred and non-preferred bar directions. Conversely, inactivating AL and PM reduced the responses of V1 neurons to their preferred orientation significantly more than (weaker) responses to non-preferred orientations in terms of absolute difference (Fig. 6A), thus indicating a non-additive form of modulation. Similarly, in terms of relative amplification (division between responses with or without optogenetic inactivation of AL or PM), responses to preferred orientations were reduced less than smaller responses to non-preferred orientations (Fig. 6B). Next, we computed the distribution of firing rate responses during either control or optogenetic inactivation (jointly for both areas), and compared these distributions to those that were obtained by either an additive or multiplicative gain modulation (Fig. 6C,D, see also STAR Methods). To achieve this, we used the average response modulation calculated considering either an additive or multiplicative gain modulation model. Compared to either a linear or multiplicative model, the experimentally observed modulation shifted the distribution of firing rates responses more strongly towards low responses (Fig. 6C,D). Altogether, these results show that AL and PM provide a form of gain modulation that selectively enhances weak responses of V1 neurons, i.e. those evoked by bars moving along non-preferred orientations and directions. In contrast, under anesthesia we found that the effect of AL and PM inactivation on the activity of V1 was compatible with a multiplicative gain (Fig. S5) - notice the relatively flat line for the response amplification plots of Fig. S5B. To further explore the mechanism underlying this form of gain modulation, and in particular whether AL and PM targeted specific neuronal subpopulations in V1, we subdivided neurons based on whether their action potential waveform was broad or narrow, i.e. corresponding to a putative pyramidal or fast-spiking interneuron, respectively (Olcese et al., 2013, 2016; Vinck et al., 2015), and based on whether neurons were located in supragranular, granular or infragranular layers (see Fig. S6A-C and STAR Methods). However, no difference was observed between putative excitatory and inhibitory neurons, nor between putative excitatory neurons residing in different cortical layers as concerns changes in visual responses following inactivation of AL and PM (Fig. S6D-G). Therefore, the modulation provided by AL and PM onto V1 seems to similarly affect all major neuronal components of V1.

**Figure 6.**
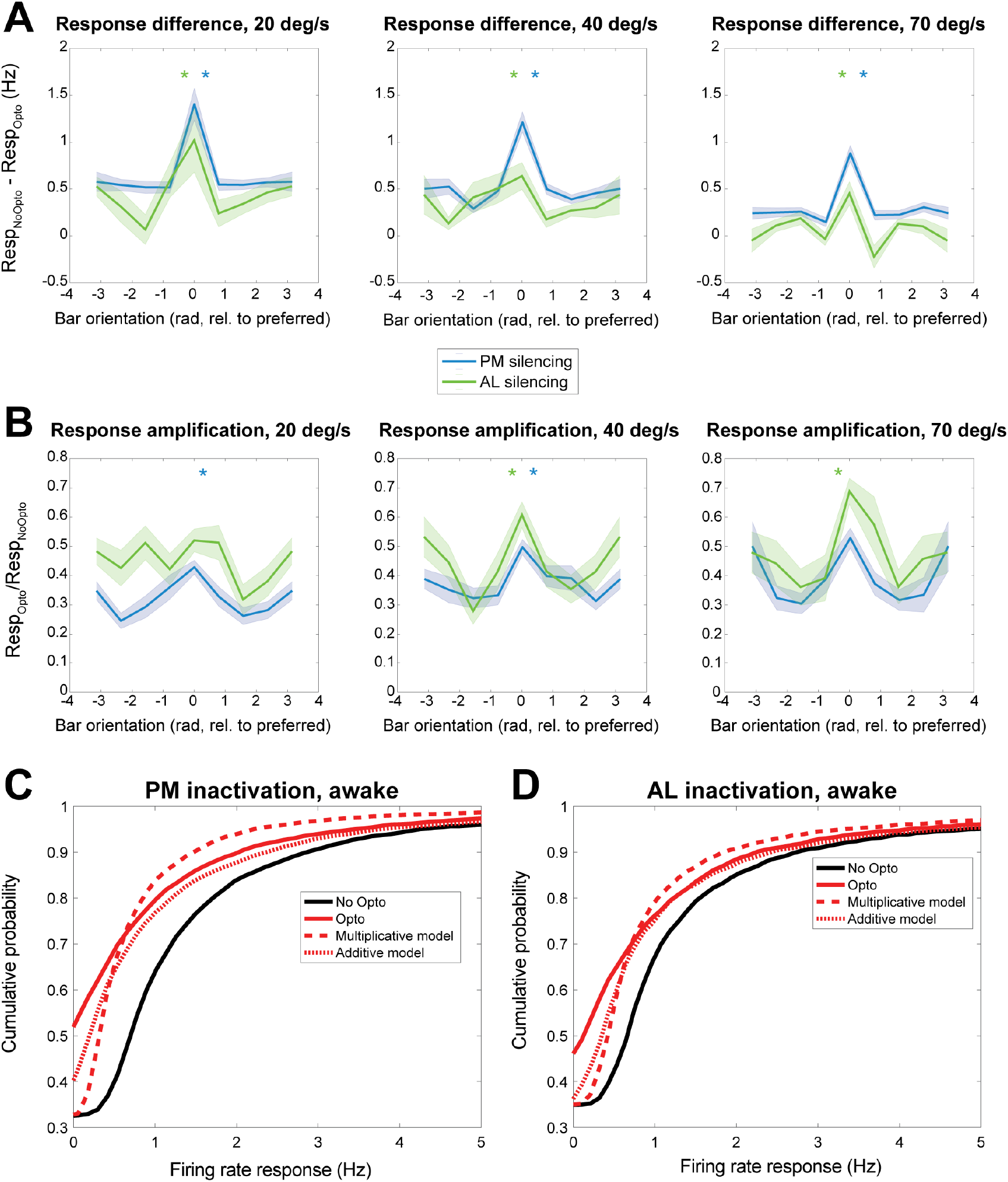
Areas AL and PM enhance visual responses in V1 during wakefulness neither by additive nor multiplicative gain. **A.** Difference in sensory-evoked responses of V1 neurons in the absence or presence of optogenetic inactivation of either area PM (blue lines) or AL (green lines) during awake recordings, separately for each orientation of a moving bar - respectively - at 20 deg/s (left), 40 deg/s (middle) or 70 deg/s (right). Curves with shading indicate mean ± SEM. The asterisks indicate if, separately for each bar speed and area (as indicated by color of asterisk), the difference in sensory evoked responses was the same for all orientations (null hypothesis, corresponding to a flat line in the plot), or not (p<0.05, one way anova; a post-hoc Tukey test revealed a larger difference for the preferred orientation corresponding to 0 rad). **B.** Same as *A*, but now for the ratio between sensory-evoked responses of V1 neurons in the presence or absence of optogenetic inactivation of either area PM (blue lines) or area AL (green lines). In general, responses to the preferred orientation were reduced less by optogenetic inactivation of area PM or AL, compared to responses to non-preferred orientations (p<0.05, oneway Anova with post-hoc Tukey test). **C.** Cumulative distribution of sensory evoked responses in V1 in the absence (black) or presence (red, solid line) of optogenetic inactivation of area PM during awake recordings. The distribution of firing rates in the presence of optogenetic inactivation of area PM was compared to that obtained when applying an additive (red, dotted line) or a multiplicative (red, dashed line) model - see STAR Methods for details. Compared to both models, optogenetic inactivation more strongly enhanced the proportion of low firing rates. **D.** Same as *E*, but now for the inactivation of area AL. See also Fig. S5 and S6.

## Discussion

In this study we investigated how two HVAs, areas AL and PM, shape visual responses in V1. Our results indicate that both areas play a similar role in amplifying responses to non-preferred stimuli in V1 as well as in enlarging receptive fields. This amplification is such that strong responses of V1 neurons to visual stimuli (e.g., bars moving along the preferred orientation of a neuron) are enhanced relatively less than weak responses (i.e. to bars moving along non-preferred orientations; Fig. 2, 6). This modulation of V1 activity, besides enhancing sensory-evoked responses non-selectively, has a seemingly detrimental effect on feature tuning, as it decreased orientation and direction selectivity of V1, both at the single-neuron and population level (Fig. 4, 5). Finally, AL and PM activity similarly affects all cortical layers and the major neuronal subpopulations in V1, and the effects are similar in wakefulness and isoflurane anesthesia.

### Areas AL and PM impact mostly on non-preferred stimulus processing in V1

Inactivation of either area AL or PM similarly reduced sensory-evoked responses in V1 (Fig. 2). Therefore, the main role of AL and PM, in our experimental paradigm, is to enhance sensory-evoked responses in V1, in particular those to non-preferred stimuli (Fig. 6). Previous studies suggested that inactivation of HVAs (Pafundo et al., 2016) or feedback projections from HVAs onto V1 (Huh et al., 2018) decrease sensory-evoked responses in V1. However, in previous studies the effects were either limited to responses to the preferred orientation (Pafundo et al., 2016), were specific for the receptive field center and not the surround (Nurminen et al., 2018), or were functionally specific (i.e. related to the tuning properties of individual HVAs). For instance, Huh et al. (2018) reported that inactivation of feedback projections from AL or PM onto V1 specifically reduced responsiveness of V1 neurons tuned to spatial frequencies similar to those of AL or PM. In contrast with that study, we observed a generalized decrease in V1 responses that was independent of the speed tuning of either AL, PM, or individual V1 neurons. In line with this reduction in sensory-evoked responses, we also observed that inactivation of AL and PM reduced receptive field size in V1. This functionally aspecific effect is even more unexpected when considering that anatomical projections from HVAs onto V1 are also functionally specific (Marques et al., 2018) and target patches in layer 1 of V1 (D’Souza et al., 2019; Ji et al., 2015) based on factors such as orientation/direction tuning and speed preference. Thus, although the response properties of HVAs as well as their connections from and to V1 are all functionally specific (Andermann et al., 2011; D’Souza et al., 2019; Federer et al., 2020; Glickfeld et al., 2013; Huh et al., 2018; Kim et al., 2018; Marques et al., 2018; Marshel et al., 2011), we found that areas AL and PM also implement a generalized, aspecific gain modulation of V1 activity. How are these two apparently discordant effects compatible?

Several factors could explain this apparent discrepancy. First, not all previous studies focused on areas AL and PM, but rather on the lateromedial (LM) secondary visual area (Marques et al., 2018; Pafundo et al., 2016). While areas such as LM, AL and PM are all considered HVAs, LM is thought to be part of the mouse ventral stream, while both AL and PM are attributed to the dorsal stream (Wang et al., 2012). Thus, the roles of area LM versus AL and PM in modulating activity in V1 might be different, also in view of the specific functions of LM in higher order visual processing, such as illusory contour detection and shape categorization (Khastkhodaei et al., 2016; Matteucci et al., 2019; Pak et al., 2020; Tafazoli et al., 2017). Previous studies indicated that inactivation of LM primarily affects the superficial layers of V1 (Pafundo et al., 2016); conversely, we observed that inactivation of AL or PM elicited a modulation that was similar for all the major subpopulations of V1. None of the prior studies that focused on AL and PM (and in general no study on HVAs) used moving bars, as the most commonly used visual stimuli were drifting gratings or Gabor patches. The use of gratings instead of bars might explain all observed differences between our and previous studies. Gratings are commonly displayed over a larger field of view (often the full visual field) compared to bars (which, in our case, were only 3 deg wide stimuli over a grey background). Thus, moving gratings simultaneously evoke activity in a larger population of neurons compared to bars, which instead activate sparser subsets of neurons sequentially, based on where in the visual field the bar is located at any moment. Therefore, at the population level, moving bars elicit an overall instantaneous weaker activity compared to gratings. The aspecific modulatory effect we report primarily affects responses to nonpreferred stimuli, but it is possible that it might occur jointly with a functionally specific form of modulation (i.e. dependent on the spatial and temporal tuning properties of HVAs and V1 neurons), such as that described for instance in Huh et al. (2018). Indeed, when we performed preliminary experiments with drifting gratings, we found results in line with (Huh et al., 2018). Specifically, we confirmed that inactivating PM during wakefulness has a stronger effect than AL on neurons preferentially responding to gratings with spatial and temporal frequencies close to those preferred by AL neurons (Fig. S1G). In line with Huh (2018), we also found a trend towards a stronger effect of silencing AL or PM the closer the speed preference of V1 neurons matched that - respectively - of PM and AL (Fig. S1G). Surprisingly, however, an opposite trend was found under anesthesia (Fig. S1H), a brain state that was not investigated by (Huh et al., 2018).

Another difference between our study and (Huh et al., 2018) is that we completely inactivated AL and PM, and not just the neurons in these areas projecting back to V1. It may be the case that inactivation of a network node, and not just of feedback-projecting neurons, may have a broader, less specific effect. While inactivating recurrently projecting neurons will primarily affect their V1 targets (but also other areas they project to), the manipulation we implemented allowed us to study the overall causal effects that AL and PM, as network nodes in the visual system, exert on V1 activity.

In view of the somewhat contrasting results from other studies using drifting gratings as visual stimuli, it will be important in future studies to compare the role of areas such as AL, PM, and also LM in modulating V1 responses to a wider array of visual stimuli. While our data suggest that indeed V1 responses to bars and gratings might be differentially affected by inactivation of AL and PM (see Fig. 2E-F and Fig. S1G), we only performed preliminary experiments to address this issue, and a complete comparison between bars and gratings is outside the scope of this study. Similarly, we primarily collected and analyzed data from V1. A deeper investigation of how areas AL and PM differentially respond to bars, gratings and other stimuli will be necessary to fully interpret their network role.

An additional contrast between our and previous studies lies in the minor differences that we observed between the effect of inactivating AL and PM during wakefulness or isoflurane anesthesia. The main difference that we found between the two brain states was a switch from non-multiplicative (and non-additive) to multiplicative gain modulation (cf. Fig. 6 and Fig. S5). Conversely, other studies reported that inactivation of HVAs more strongly affects V1 during wakefulness than anesthesia (Keller et al., 2020; Vangeneugden et al., 2019). This weaker effect of top-down modulation under anesthesia is in line with results from human subjects during loss of consciousness (Boly et al., 2011; Sikkens et al., 2019). A likely explanation for these different results lies again in the different type of stimulus that we used, which evokes less powerful activity changes in visual cortices compared to gratings and Gabor patches.

### Enhanced sensory-evoked responses in V1 come at the expense of orientation and direction selectivity

The enhancement of sensory-evoked responses that AL and PM induce in V1 comes, strikingly, at the expense of orientation and direction selectivity. A first interpretation holds that, from an evolutionary perspective, V1’s responses to any stimulus (whether preferred or non-preferred at the single-neuron level) may be more important than encoding detailed features - orientation and direction - of the stimuli. This is true both when considering single neurons, as quantified by gOSI and gDSI, as well as population-level coding. This suggests that the weaker orientation and direction tuning which is present in V1 with functionally intact AL and PM is likely sufficient to enable a proper processing of visual stimuli. It is possible, however, that different results might have been obtained with other types of visual stimuli, such as moving gratings (e.g. (Jin and Glickfeld, 2020)). Therefore, the visual system might operate in a regime that balances the processing of stimulus features such as orientation and direction with the ability to process stimuli which are smaller and (at least at the single-neuron level) less salient.

A second interpretation of our results is that HVAs such as AL and PM might provide contextual, predictive representations to V1 via recurrent projections. Following a predictive processing framework (Friston, 2005; Pennartz et al., 2019; Rao and Ballard, 1999), this contextual information may be relatively uninformative in the case of the small bars moving through an otherwise blank field, and therefore feedback from HVAs onto V1 may enhance the uncertainty and associated error information coded at the level of V1 neurons: this would explain the reduced responses to non-preferred and preferred stimuli as well as reduced receptive field size under AL or PM inactivation (Fig. 2,3). Moreover, this framework may explain why our results differ considerably from previous studies that used moving gratings, because the latter type of stimulus conveys a much higher spatial predictability across the visual field than an isolated moving bar.

Of relevance, the effects of AL and PM inactivation were similar on single-neuron orientation and direction tuning, but different in terms of population decoding: inactivating AL improved population decoding of stimulus direction more than inactivation of PM did (Fig. 5). Recent studies also identified different functions of AL and PM in orientation discrimination and spatial integration, with PM showing larger receptive fields than AL (Murgas et al., 2020) and no involvement (in contrast with AL) in orientation discrimination (Jin and Glickfeld, 2020). To improve our understanding of the different properties of AL and PM, further experiments will be required to better characterize how single-neurons and populations across V1 and distinct HVAs process information about isolated moving objects (such as bars) compared to spatially broader stimuli, which may be associated with more contextual information and thus differentially engage AL and PM (Murgas et al., 2020).

### Higher order visual areas broaden stimulus responsiveness in V1

The classical framework to interpret visual processing in the neocortex follows a hierarchical approach, in which each subsequent processing stage is tasked with processing more complex stimulus features (Felleman and Van Essen, 1991; Riesenhuber and Poggio, 1999). HVA properties seem to support this view, as some higher-order visual processing is either directly dependent upon, or facilitated by them (Khastkhodaei et al., 2016; Matteucci et al., 2019; Pak et al., 2020; Tafazoli et al., 2017). Nevertheless, recent studies have shown that some key features of early visual processing, at the stage of V1, are enabled by virtue of top-down modulation originating in HVAs (Keller et al., 2020; Vangeneugden et al., 2019). Our study supports this notion, by indicating that, beyond being involved - as previously shown - in the development of response features such as surround suppression (Vangeneugden et al., 2019) and complex receptive fields (Keller et al., 2020), HVAs are also instrumental in the sculpting of sensory-evoked responses themselves. To understand how the neocortex processes visual stimuli, it is therefore essential to look beyond individual areas, and beyond a mainly hierarchical - and feedforward - information flow. While visual information processing in V1 certainly stems from thalamo-cortical projections, our results reveal that feedback information from HVAs also contributes to basic properties of V1 such as responses to oriented bars, in addition to being involved in the generation higher-level representations - e.g. of illusory contours (Pak et al., 2020). Our experiments showed that AL and PM enhance responses to non-preferred stimuli in V1 more than responses to preferred stimuli. This effect may be explained by an added level of background excitation that HVAs could provide to V1 neurons. Such additional excitation might modify the supposedly sigmoid input-output transfer function of V1 neurons in a way that more strongly amplifies weak inputs compared to strong ones. This preferential enhancement of weaker sensory-evoked responses is reminiscent of the principle of inverse effectiveness, which is commonly observed in multisensory integration, both in subcortical (Stanford et al., 2005; Stein and Stanford, 2008) and cortical regions (Meijer et al., 2019, 2020; Olcese et al., 2013). In multisensory integration regions, responses to weak unisensory stimuli occurring simultaneously are amplified - in relative terms - more than responses to strong unisensory stimuli. Such a phenomenon may be advantageous for the detection of weak sensory stimuli (Stein and Stanford, 2008). By broadening stimulus responsiveness, in particular to non-preferred and small visual features, HVAs such as AL and PM might play a role akin to that fulfilled by inverse effectiveness in aiding the detection of visual stimuli which would otherwise not evoke a significant response. This may provide a behavioral advantage by enabling to process (and thus detect), small, barely noticeable visual stimuli.

Finally, it must be pointed out that our experiments were performed in awake, but passive animals. Therefore, in future studies, it will be essential to study the structure and function of feedback from HVAs in animals trained in visual detection or discrimination to understand the functional role of feedback projections in attention, predictive representation, detection and discrimination (Gilbert and Li, 2013; Pennartz et al., 2019; Rao and Ballard, 1999; Zhang et al., 2014).

### Conclusions

Higher order visual areas are key elements of the cortical network of visual processing, as they play a role not only in further analyses of visual information after V1, but also in modulating V1 activity itself. Here we show how two HVAs with different response properties (AL and PM) similarly enhance responses of V1, especially to non-preferred visual stimuli. At the population level, AL and PM activity makes it easier for V1 to respond to moving bars, but at the same time more difficult to decode their precise orientation and direction. Areas AL and PM therefore provide a major contribution to sculpting of V1 responses to simple visual objects: they effectively broaden sensory tuning curves and thus render neurons more responsive to a broad range of orientations.

## Acknowledgements

This work was supported by the European Union’s Horizon 2020 Framework Program for Research and Innovation under the Specific Grant Agreement 720270 (Human Brain Project SGA1) to C.M.A.P., Grant Agreement 785907 (Human Brain Project SGA2) and 945539 (Human Brain Project SGA3) to C.M.A.P. and U.O., by the FLAG-ERA JTC 2015 project CANON (co-financed by the Netherlands Organization for Scientific Research - NWO) to U.O. and by the FLAG-ERA JTC 2019 project DOMINO (co-financed by NWO) to U.O. The authors would like to thank Laura Bavelaar for her support in the initial phases of this project.

## Author Contributions

Conceptualization, U.O.; Methodology, U.O. and M.O.L.; Software, M.O.L., U.O. and A.C.C; Formal analysis, M.O.L. and U.O., Investigation, M.O.L., L.B. and U.O.; Data Curation, M.O.L.; Writing - Original Draft, U.O. and M.O.L.; Writing - Review & Editing, U.O., M.O.L. and C.M.A.; Visualization, U.O.; Supervision, U.O. and C.M.A.; Funding Acquisition, C.M.A. and U.O.

## STAR Methods

### KEY RESOURCE TABLE

| REAGENT or RESOURCE | SOURCE | IDENTIFIER |
| --- | --- | --- |
| **Bacterial and Virus Strains** |  |  |
| AAV1/5-EF1a-double floxed-hChR2(H134R)-EYFP-WPRE-HGHpA | AddGene | 20298-AAV1 |
| **Chemicals, Peptides, and Recombinant Proteins** |  |  |
| DiI | ThermoFisher Scientific | Catalog number: V22889 |
| **Experimental Models: Organisms/Strains** |  |  |
| Mouse: PV<sup>cre</sup> (B6;129P2-Pvalbtm1(cre)Arbr/J | Jackson Laboratory (mice bred in own facility) | 008069 |
| Mouse: Parvalbumin-TdTomato C57BL/6-Tg(Pvalb-tdTomato)15Gfng/J | Jackson Laboratory (mice bred in own facility) | 027395 |
| **Laminar probes for electrophysiological recordings** |  |  |
| 32 channel, chronic Poly2 probe | Neuronexus | A1x32-Poly2-10mm-50s-177 |
| 32 channel, chronic 4-shank probe | Neuronexus | A4x8-5mm-100-200-177 |
| 64 channel, acute Poly2 probe | Neuronexus | A1x64-Poly2-6mm-23s-160 |
| **Software and Algorithms** |  |  |
| MATLAB | Mathworks | https://www.mathworks.com |
| Python | Python | https://www.python.org |
| R | The R Foundation | https://www.r-project.org/ |
| Klusta | Rossant et al., 2016 | https://klusta.readthedocs.io/ |
| Phy Kwik-GUI | Rossant et al., 2016 | https://phy.readthedocs.io/ |
| **Other** |  |  |

### CONTACT FOR REAGENT AND RESOURCE SHARING

This study did not generate new unique reagents. Any request about resource sharing should be directed to Umberto Olcese or Matthijs oude Lohuis.

### EXPERIMENTAL MODEL AND SUBJECT DETAILS

#### Animals

All animal experiments were performed according to the national and institutional regulations. The experimental protocol was approved by the Dutch Commission for Animal Experiments and by the Animal Welfare Body of the University of Amsterdam. A total of 14 male mice from two transgenic mouse lines were used: PVcre (B6;129P2-Pvalbtm1(cre)Arbr/J, JAX mouse number 008069) and PVcre/TdTomato (C57BL/6-Tg(Pvalb-tdTomato)15Gfng/J, JAX mouse number 027395). Animals were at least 8 weeks of age at the start of experiments. Mice were group housed, with ad libitum access to water and food, under a reversed day-night schedule (lights were switched off at 8:00 and back on at 20:00). All experimental procedures were performed during the dark period.

### METHOD DETAILS

#### Headbar implantation

Mice were subcutaneously injected with the analgesic buprenorphine (0.025 mg/kg) and maintained under isoflurane anesthesia (induction at 3%, maintenance at 1.5-2%) during surgery. The skin above the skull was epilated, disinfected, and a circular area was removed with the edges glued to the outer parts of the skull using tissue adhesive (3M Vetbond, MN, United States) to prevent post-surgical infections. A custom-made titanium head-bar with a circular recording chamber (inner diameter: 5 mm) was positioned over the exposed skull of the left hemisphere to include visual, auditory and somatosensory cortices and attached using cyanoacrylate and C&B Super-Bond (Sun Medical, Japan).

#### Intrinsic optical imaging

To localize individual higher visual cortical areas we performed intrinsic optical imaging (IOI) under lightly anesthetized conditions (0.7-1.2% isoflurane). A vasculature image was acquired under 540 nm light before starting the imaging session. During IOI, the cortex was illuminated with monochromatic 630 nm light. Images were acquired at 1 Hz using an Adimec 1000m CCD camera (1004 x 1004 pixels) connected to a frame grabber (Imager 3001, Optical Imaging Inc, Germantown, NY, USA), defocused about 500-600 μm below the pial surface. We presented visual, auditory and tactile stimuli. Visual stimuli consisted of full field drifting gratings (spatial frequency 0.05 cpd, temporal frequency 1.5 Hz) for 1 second in each of 8 directions. Auditory stimuli consisted of alternations between chirps sweeping up or down in frequency (1-40 kHz) and band-passed white-noise (1-40kHz) calibrated at 70 dB Sound Pressure Level (SPL). Tactile stimuli were full whisker-pad deflections driven by a piezo-actuator (18° angle). For each type of stimulation we acquired 8 seconds of baseline signal and 8 seconds of hemodynamic response during stimulation. The acquired frames during the response were baseline-subtracted, averaged and thresholded to produce a map of localized individual primary and higher order areas. PM and AL were identified based on the IOI signal map in combination with previously published maps (Glickfeld and Olsen, 2017; Olcese et al., 2013; Wang and Burkhalter, 2007) and marked on the skull based on the vasculature image. After IOI, the recording chamber was covered with silicon elastomer (Picodent Twinsil) and mice were allowed to recover for 2-7 days.

#### Viral injections

Mice were subcutaneously injected with the analgesic buprenorphine (0.025 mg/kg) and maintained under isoflurane anesthesia (induction at 3%, maintenance at 1.5-2%) during surgery. We performed a small craniotomy over the area of interest (either PM or AL, identified using IOI) using a dental drill and inserted a glass pipette backfilled with AAV2.1-EF1a-double floxed-hChR2(H134R)-EYFP-WPRE-HGHpA (titer: 7×10^12^ vg/ml, 20298-AAV1, Addgene). Four injections of 13.8 nl were made at two depths (two at 700 μm and two at 400 μm below the dura) using a Nanoject pressure injection system (Drummond Scientific Company, USA). Each injection was spaced apart by at least 5 minutes from the next one to promote diffusion and prevent backflow. After viral injections, the recording chamber was covered with silicon elastomer (Picodent Twinsil) and mice were allowed to recover.

#### Craniotomy

After at least 3 weeks to allow for robust viral expression, mice were subcutaneously injected with the analgesic buprenorphine (0.025 mg/kg) and maintained under isoflurane anesthesia (induction at 3%, maintenance at 1.5–2%) during surgery. We performed small (200 um) craniotomies over the areas of interest (V1 and either PM or AL) using a dental drill. The dura was left intact if possible. The recording chamber was sealed off with silicon elastomer and the mice were allowed to recover for 24h.

#### In vivo electrophysiology

Mice were fixated in a custom-built holder in a dark and sound-attenuated cabinet. The body of the mouse was put in a tube (diameter: 4 cm) to limit body movements. The headbar was attached to a custom-made holder via two screws. Before recording sessions, mice were habituated to this type of head-fixation by daily progressive incremental time spent in head-fixation.

Recordings were performed either in an awake or anesthetized state and the order was counterbalanced across recording days. Under anesthesia, pure oxygen with isoflurane (at 0.6-1.2%) was delivered at 0.8 l/min. The level of anesthesia was monitored by observing breathing rate and neural activity. Isoflurane levels were slowly lowered over the course of a recording session to counteract tissue build-up and maintain a stable depth of anesthesia. Body temperature was monitored throughout and kept at 37.5 °C.

Extracellular recordings were performed with 32- or 64-channel microelectrode arrays (NeuroNexus, Ann Arbor, MI-A1×32-Poly2-10mm-50s-177, A4×8-5mm-100-200-177, or A1×64-Poly2-6mm-23s-160). Each recording session the electrode arrays were slowly inserted until the recording sites spanned the cortical layers. We verified visual responsiveness by displaying full-field gratings and reinserted the electrodes if there was no robust visual responsiveness in neural activity. The number of recording sessions was limited to 3 to minimize recording from a damaged circuit. For some recording sessions electrodes were dipped in DiI (ThermoFisher Scientific) allowing better post hoc visualization of the electrode tract (Fig. 1C). After insertion, the exposed cortex and skull were covered with 1.3-1.5% agarose in artificial CSF to prevent drying and to help maintain mechanical stability. The ground was connected to the headbar and the reference electrode to the agarose solution. Recordings started at least 15 minutes after insertion to allow for tissue stabilization. Neurophysiological signals were amplified (x1000), bandpass filtered (0.1 Hz to 9 kHz) and acquired continuously at 32 kHz with a Digital Lynx 128 channel system (Neuralynx, Bozeman, MT).

#### Optogenetics

To locally photostimulate PM or AL, a 473 nm laser (Eksma Optics, Vilnius, Lithuania, DPSS 473nm H300) was connected with a fiber-optic patch cord to a fiber-optic cannula (ID 200 um, NA 0.48, DORIC lenses) that was positioned directly over the thinned skull at the area of interest. Photostimulation consisted of 10 ms pulses delivered at 20 Hz for the duration of visual stimulus presentation. Stimulus duration varied depending on the traversal time of the bar across the screen and, depending on travelled distance and speed of the bar, ranged from 0.45 s (vertical bar moving at 70°/s) to 6.3 s (diagonal bar moving at 20°/s). Light delivery was controlled by a shutter (Vincent Associates LS6 Uniblitz). During each session we simultaneously performed extracellular recordings in the areas of interest (V1 and either PM or AL) and adjusted laser power to the minimum that maximally inhibited neural activity (routinely 5-20 mW total power).

#### Visual stimulation

Visual stimuli were gamma-corrected and presented with a 60 Hz refresh rate on an 18.5 inch monitor positioned at a 45° angle with the body axis from the mouse at 21 cm from the eyes, subtending 91° horizontally and 60° vertically. Three sets of visual stimuli were used.

##### Checkerboards

Before each session, we displayed full-field contrast-reversing checkerboards (full contrast, spatial frequency = 10 retinal degrees, temporal frequency of contrast reversal = 0.5 Hz, n=10 reversals) to estimate laminar electrode positioning (see below).

##### Bars

Each bar stimulus consisted of a single white bar drifting across an isoluminant gray screen in one of 8 directions at one of three speeds (20°/s, 40°/s or 70°/s) either in absence or presence of photostimulation. Stimuli were separated by an inter-trial interval of 3 seconds and repeated 20 times. The total trial set therefore consisted of 8 (orientations) x 3 (speeds) x 2 (photostimulation conditions) x 20 (repetitions) = 960 trials.

##### Gratings

Grating stimuli consisted of full-field drifting square-wave gratings (70% contrast) for 2 seconds, separated by 2 seconds inter-trial interval. Similar to the bar stimuli, gratings drifted in one of 8 directions at one of three speeds (20°/s, 40°/s or 70°/s) either in absence or presence of photostimulation for 20 repetitions. The three speeds were constructed based on combinations of spatial and temporal frequencies to optimize V1, PM and AL responsiveness (Andermann et al., 2011; Marshel et al., 2011): Slow 20°/s: Spatial frequency = 0.1 cpd, Temporal frequency = 2 Hz, Mid 40°/s: Spatial frequency = 0.075 cpd, Temporal frequency = 3 Hz, Fast 70°/s: Spatial frequency = 0.057 cpd, Temporal frequency = 4 Hz.

#### Histology

At the end of the experiment, mice were overdosed with pentobarbital and perfused with 4% paraformaldehyde in phosphate-buffered saline, and their brains were recovered for histology. We cut coronal 50 μm sections with a vibratome, stained them with DAPI (0.3 μM) and imaged mounted sections to verify the viral expression and recording sites. The borders of individual higher visual areas in individual animals are not definable based on an atlas. However, with this consideration in mind, data from five animals was excluded based on weak expression in putative PM or AL or strong off-target expression beyond PM or AL or into V1.

### QUANTIFICATION AND STATISTICAL ANALYSES

#### Spike sorting

Before spike detection the median of the raw trace of nearby channels (400<um) was subtracted to remove artefacts. Spike detection and sorting were done using Klusta and manual curation using the Phy GUI (Rossant et al., 2016). During manual curation each proposed single unit was inspected based on its waveform, autocorrelation function and firing pattern across channels and time. Only high-quality single units were included that (1) had an isolation distance higher than 10 (Schmitzer-Torbert et al., 2005), (2) had less than 0.1% of their spikes within the refractory period of 1.5 ms, (3) were present throughout the session.

#### Classification neuron subtypes

Putative pyramidal and putative fast-spiking interneurons were separated based on the peak-to-trough delay of their average normalized action potential waveform (Niell and Stryker, 2008). The peak-to-trough delay was computed as the time between peak positive and peak negative voltage deflection (in ms) and single units with a delay lower than 0.45 ms were classified as narrow-spiking, while units with a delay higher than 0.55 ms were classified as broad-spiking. The rest remained unclassified. In total 76.9% were labeled as broad spiking, 20.6% as narrow-spiking and 2.5% as unclassified.

#### Laminar depth estimation

The laminar depth of each electrode was estimated based on current source density analysis (CSD) of the local field potential (LFP) in response to contrast-reversing checkerboard stimuli (see above). The CSD profile was computed by applying standard Nicholson-Freeman calculations on the low-pass filtered signal (<100 Hz, 4th order Butterworth filter) with Vaknin transform (Vaknin et al., 1988) with 0.4 Siemens per meter as conductivity. We calculated the CSD profile for each of the linear arrays of electrodes on our polytrode configuration separately and then merged the profiles. The electrode with the earliest visible sink was designated as the center of layer IV. Single units recorded from electrodes spanning 150 μm around this electrode were labeled as granular and units recorded from electrodes below and above this layer were labeled infra- and supragranular, respectively.

#### Firing rate response

To compute firing rates in response to visual stimuli, spikes times were aligned to stimulus onset, binned in 1 ms bins and convolved with a Gaussian window (50 ms standard deviation). To compute single trial responses for bar stimuli we first identified the time of peak response for each condition (orientation x speed) by averaging across trial repetitions without photostimulation. The response on each trial was obtained by averaging the single trial firing rate over 300 ms around this peak time (± 150 ms). For grating stimuli the single-trial firing rate was averaged over 0-1000 ms after stimulus onset. Only neurons showing a significant sensory evoked response (defined as having an average z-scored response >1 for at least one bar direction) were retained for further analyses. The following table summarizes the number of V1 neurons that were retained for analysis:

| Bar speed | 20 deg/s | 40 deg/s | 70 deg/s |
| --- | --- | --- | --- |
| PM inactivation – awake recordings | 207 | 206 | 185 |
| AL inactivation – awake recordings | 94 | 97 | 79 |
| PM inactivation – anesthetized recordings | 88 | 87 | 87 |
| AL inactivation – anesthetized recordings | 67 | 65 | 63 |

#### Quantification of tuning curves and receptive field size

Tuning curves for single neurons were quantified after computing, independently for each direction and separately for each speed, the firing rate response to a visual stimulus. The preferred orientation/direction was computed, separately for each speed, based on the bar direction eliciting the largest average firing rate response, in the absence of optogenetic inactivation. To align tuning curves, the preferred orientation of each neuron in the absence of optogenetic inactivation was set to 0 degrees and other orientations were displayed relative to this.

Receptive field size was computed separately for the average responses to each bar direction. We computed the response onset as the first time point after stimulus onset in which the z-scored firing rate response exceeded 1. The response offset was defined as the first time point following response onset for which the z-scored firing rate response dropped below 1. The receptive field size for a given direction was computed as the duration of the response (time lag between response onset and offset) multiplied by the speed of the bar. Receptive field size was aligned to the preferred direction, as described for the tuning curves.

#### Orientation and direction selectivity

Orientation and direction selectivity were computed using a global orientation selectivity index (gOSI) and a global direction selectivity index (gDSI) (Ibrahim et al., 2016; Mariño et al., 2005). These two measures were computed as:

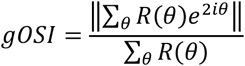

and

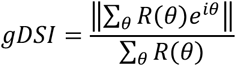

Here *R(θ)* is the firing rate response of a neuron to a bar moving along direction *θ* and *i* is the imaginary unit. gOSI and gDSI vary between 0 and 1, with 0 indicating a neuron completely untuned for orientation/direction, and 1 a neuron only responding to a single orientation/direction, respectively.

#### Decoding analysis

The population decoding analysis was done using a pseudo-population approach. Decoding was separately performed for awake and anesthetized recordings. All recorded neurons were pooled together (even if they were recorded in different sessions) and decoding was performed on a randomly selected number of neurons equal to the lowest available number of neurons per condition. For awake recordings, this amounted to 79 neurons; for anesthetized recordings, to 63 neurons (see the table above). In detail, we used the same number of neurons to decode the direction of a moving bar presented at every speed, without optogenetics, or with inactivation of either PM or AL; this procedure allowed us to fairly compare the different conditions (area being inactivated and bar speed). When pooling together data from different recording sessions, we only considered conditions (bars moving along a certain direction and speed) which had been repeated over at least 10 trials. For all conditions, 20 trials were sampled over recording sessions (with replacement, if fewer than 20 trials were present, and without replacement otherwise). This data was used to train a k-nearest neighbors classifier, which was trained to decode the direction of the bar being presented, based on the single-trial firing rate response (computed as described above as the average firing rate in a 300 ms window centered around the peak latency of the response of each neuron to a bar with a certain speed and direction). The performance of the decoder was assessed with a leave-one-out crossvalidation procedure. Training was repeated 100 times, each time with a different, random set of neurons and randomly sampled trials. For each training set, we computed the average decoding error.

#### Statistical analysis

All statistical analyses were done using parametric methods (t-tests and ANOVAs), as the assumption of normality was not violated in any instance. If applicable (i.e. when an Anova was performed), multiple comparisons were corrected using a Tukey post-hoc test. When multiple, independent comparisons were performed, p-values were corrected via the application of a Benjamini-Hockberg false discovery rate (FDR) procedure (Korthauer et al., 2019).

## DATA AND SOFTWARE AVAILABILITY

Original data and the MATLAB, Python and R scripts used to perform the analyses presented in this manuscript are available by reasonable request to Umberto Olcese.

## Supplemental Information

### Supplemental Figure Legends

**Figure S1.**
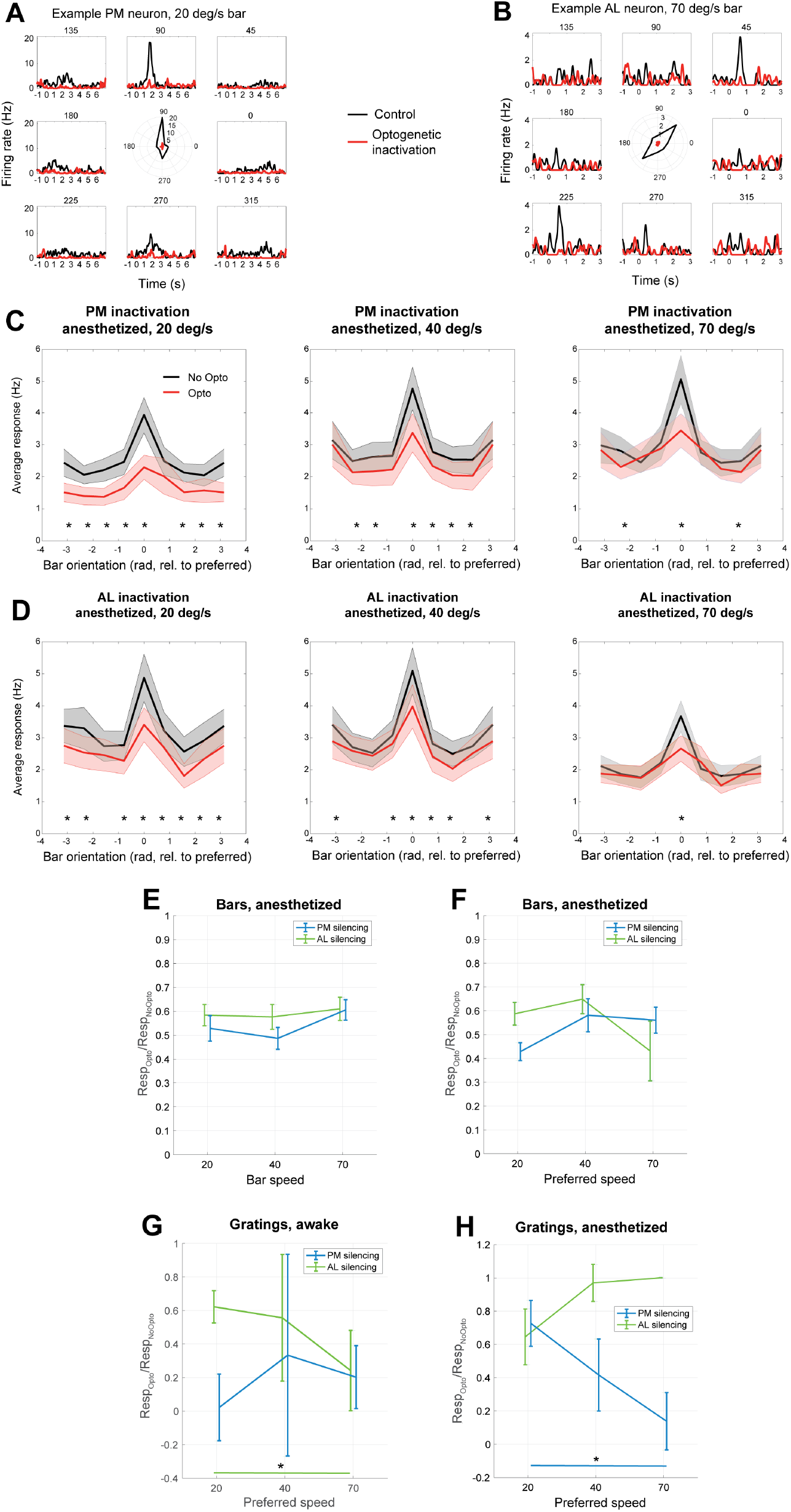
Optogenetic inactivation of area AL and PM under anesthesia depresses sensory-evoked responses in V1; experiments performed with drifting gratings instead of bars confirm the previously reported functionally-specifíc effect of inactivating AL and PM – refers to Fig. 1 and 2. **A.**Same as Fig. 1J, but now for a neuron located in area PM and responding to a bar moving at 20 deg/s during wakefulness. Black: control; red: inactivation of area PM. **B.** Same as *A*, but now for a neuron located in area AL and responding to a bar moving at 70 deg/s during wakefulness. Note how optogenetic stimulation suppresses the responses of neurons located in PM and AL (panels *A* and *B*, respectively), but only reduces the responses of the neuron located in V1 (Fig. 1J). **C.** Same as Fig. 2B, but now for recordings performed under anesthesia. **D.** Same as Fig. 2D, but now for recordings performed under anesthesia. **E.** Same as Fig. 2E, but now for recordings performed under anesthesia. **F.** Same as Fig. 2F, but now for recordings performed under anesthesia. **G.** Same as Fig. 2F, but now for responses of V1 neurons to moving gratings during awake recordings. **H.** Same as *G*, but now for recordings performed under anesthesia.

**Figure S2.**
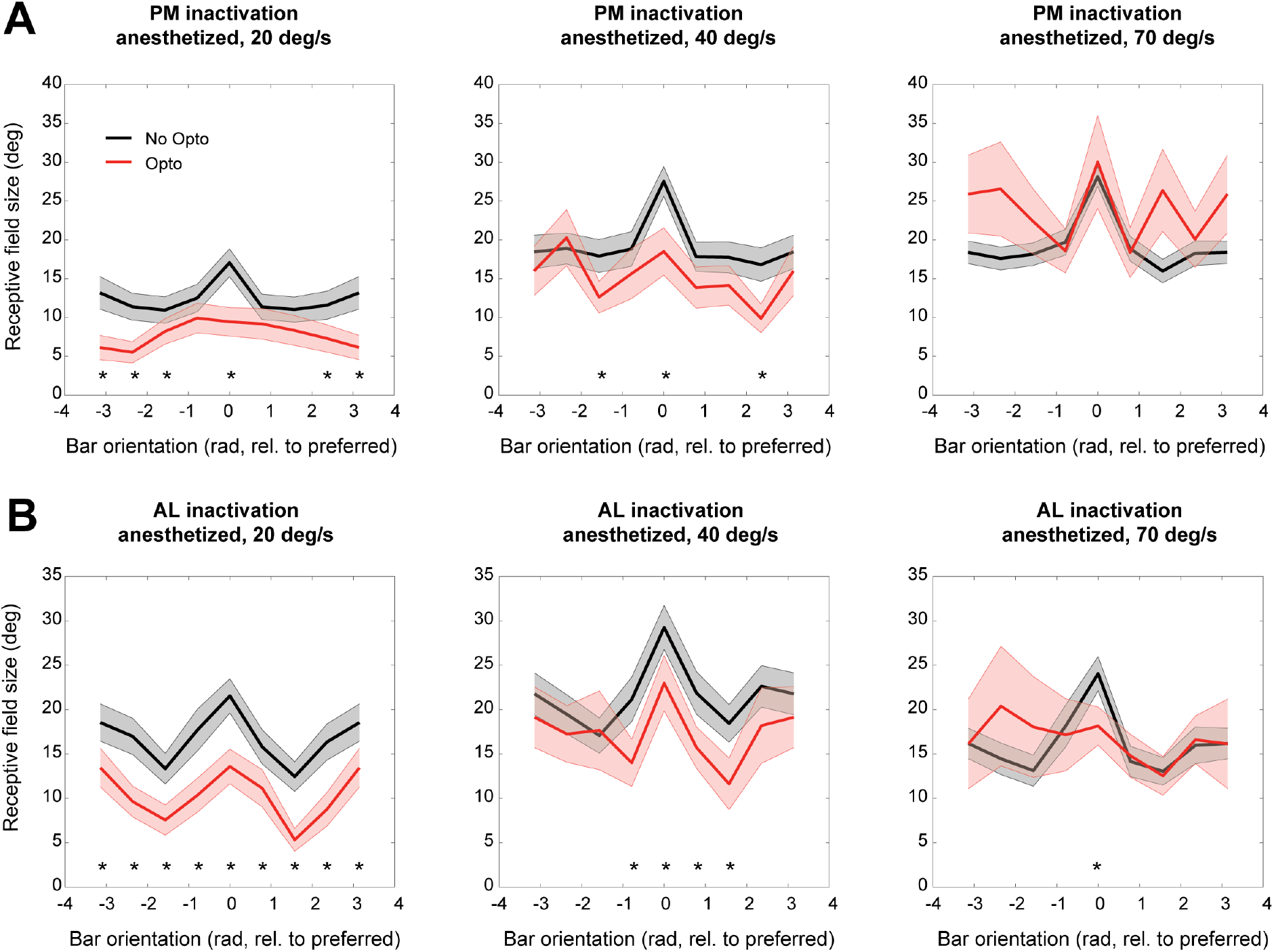
Optogenetic inactivation of area AL and PM under anesthesia reduces receptive field size of V1 neurons – refers to Fig. 3. **A.** Same as Fig. 3A, but now for recordings performed under anesthesia. **B.** Same as Fig. 3B, but now for recordings performed under anesthesia.

**Figure S3.**
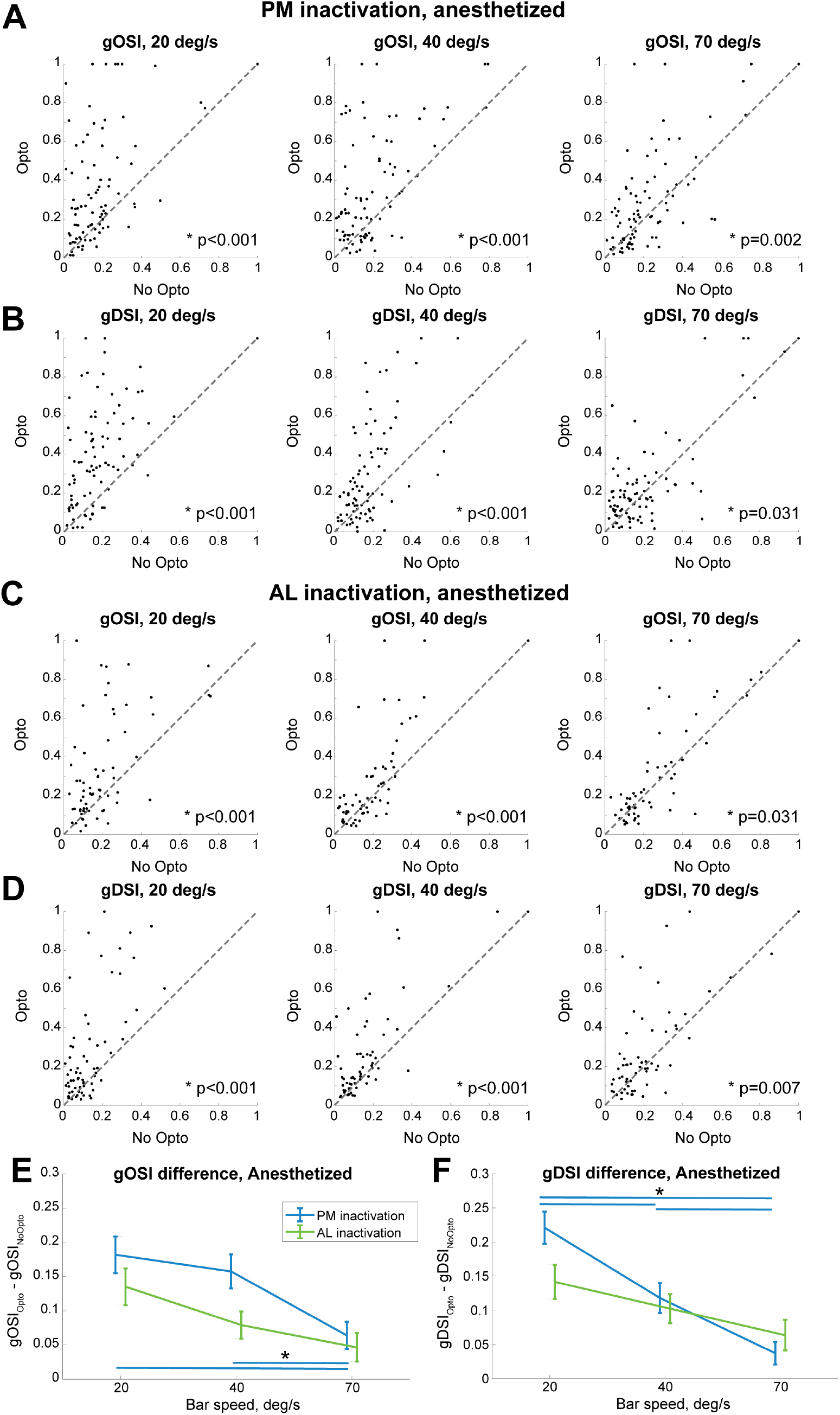
Optogenetic inactivation of area AL and PM under anesthesia enhances orientation and direction selectivity in V1 – refers to Fig. 4. **A.** Same as Fig. 4A, but now for recordings performed under anesthesia. **B.** Same as Fig. 4B, but now for recordings performed under anesthesia. **C.** Same as Fig. 4C, but now for recordings performed under anesthesia. **D.** Same as Fig. 4D, but now for recordings performed under anesthesia. **E.** Same as Fig. 4E, but now for recordings performed under anesthesia. Asterisks indicate significant differences either between speeds (for a given area) or between inactivation of distinct areas, given the same speed (p<0.05, 2-way anova with post-hoc Tukey test). **F.** Same as Fig. 4F, but now for recordings performed under anesthesia. Asterisks indicate significant difference either between speeds (for a given area), or between inactivation of distinct areas, given the same speed (p<0.05, 2-way anova with post-hoc Tukey test). In neither panel *E* nor *F* were significant differences between areas (separately for each speed) found.

**Figure S4.**
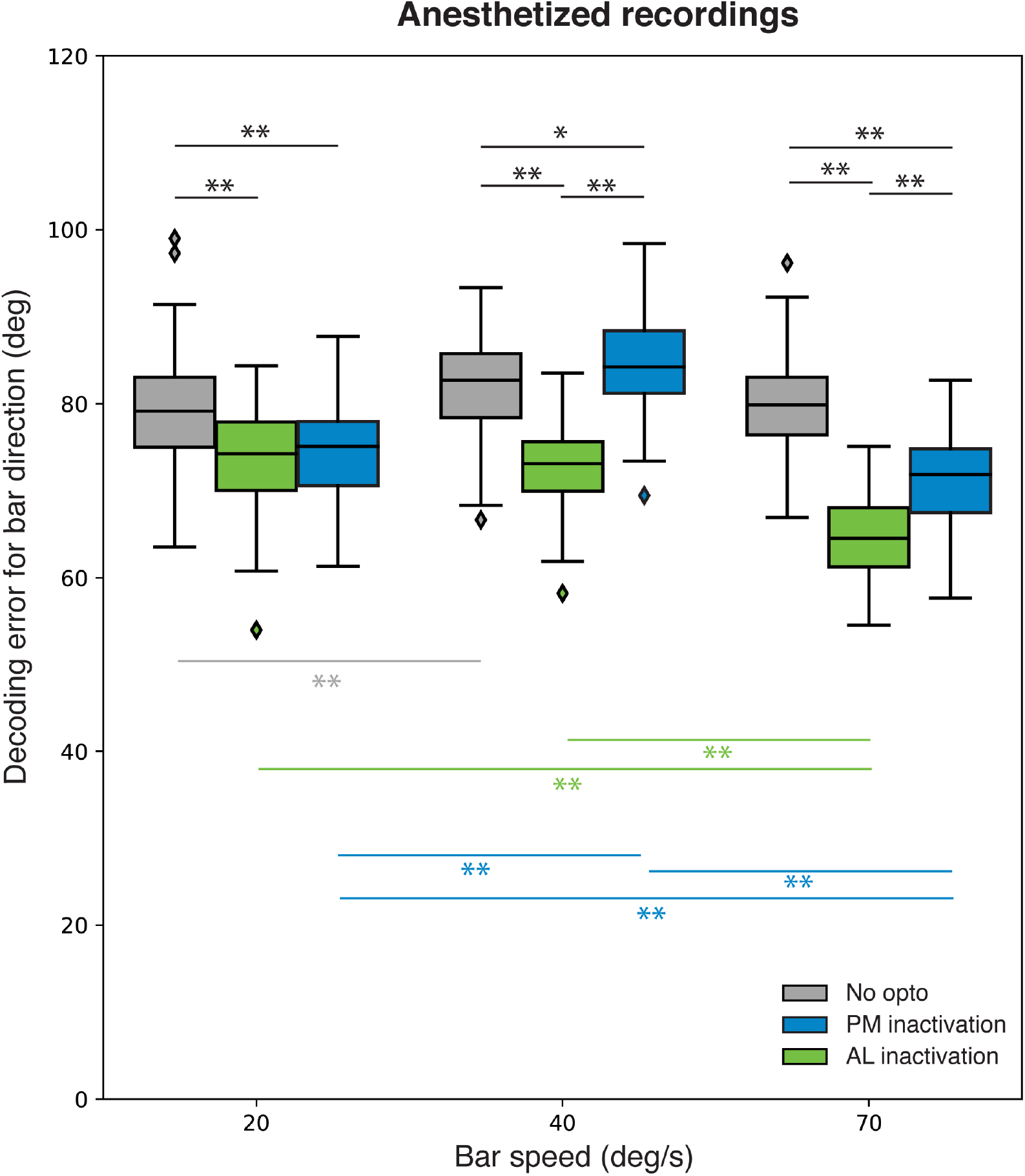
Optogenetic inactivation of area AL and PM under anesthesia enhances single-trial decoding of bar orientation in V1– refers to Fig. 5. Same as Fig. 5, but now for recordings performed under anesthesia.

**Figure S5.**
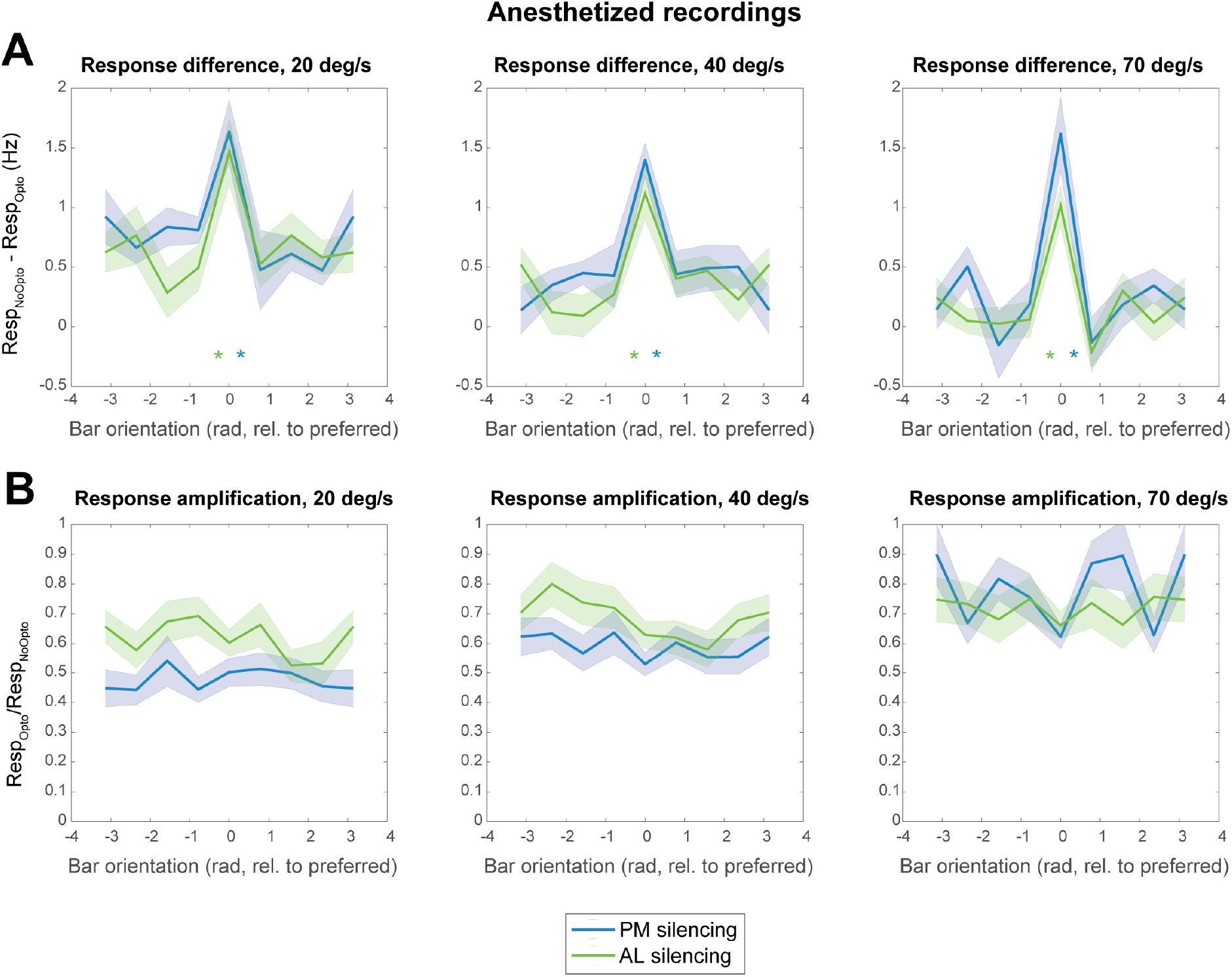
Areas AL and PM enhance visual responses in V1 under anesthesia in accordance with a multiplicative gain modulation. – refers to Fig. 6. **A.**Same as Fig. 6A, but now for recordings performed under anesthesia. **B.** Same as Fig. 6B, but now for recordings performed under anesthesia. Note the relatively flat line in all subplots, indicating that a multiplicative gain modulation applies to all orientations.

**Figure S6.**
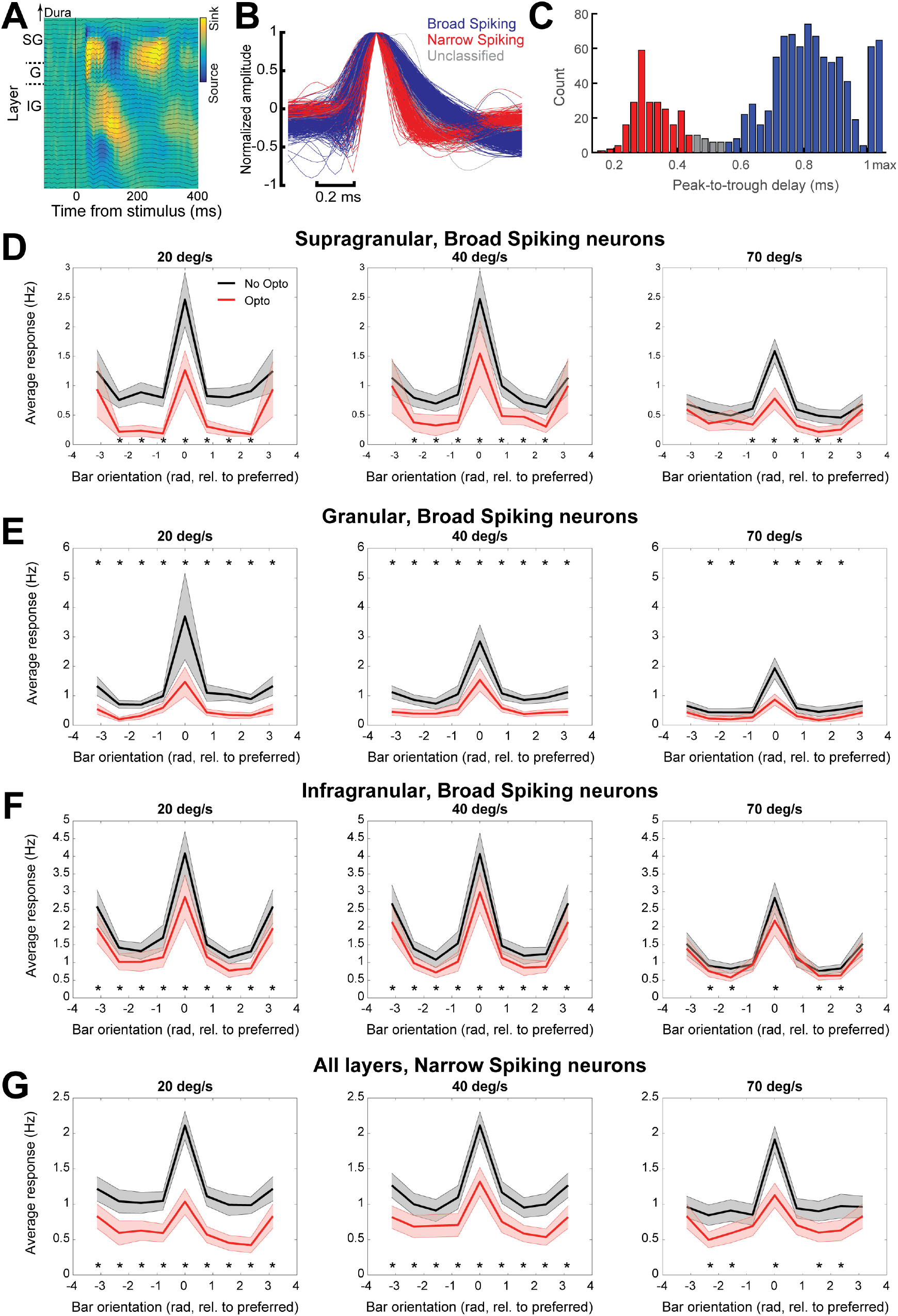
Optogenetic inactivation of area AL and PM during wakefulness similarly affects distinct V1 neuronal subpopulations– refers to Fig. 6. **A.** Current source density profile of average response to checkerboard stimulation across layers of V1. The bottom of the earliest sink after checkerboard onset was used to demarcate the boundary between L4 and L5 - as in (Schnabel et al., 2018). We used this in combination with registered depth of penetration of the silicon probe relative to the cortical surface, to align electrode depth across recordings sessions. Black traces show the local field potential traces of each channel along the electrode tract. **B.**Normalized waveforms for each individually recorded V1 neuron, averaged over all recorded action potentials and colored by their classification based on peak-to-trough delay (blue: broad spiking neuron; red: narrow spiking neuron; grey: undetermined). **C.** Histogram of peak-to-trough delay for the three classes. Bars at maximum indicate neurons whose trough extended beyond the sampled time around the action potential (2 ms). **D.** Tuning curves of average responses of broad spiking V1 neurons located in supragranular layers during awake recordings to bars moving at different orientation (aligned by the preferred orientation, which is set at 0 deg) in the absence or presence of optogenetic inactivation of either area PM or AL (black and red curves, respectively). Curves with shading indicate mean ± SEM. Asterisks indicate significant differences between responses to bars moving at a given orientation in the absence or presence of optogenetic inactivation of area PM (p<0.05, paired t-test, FDR-corrected). From left to right: responses to bars moving at 20, 40 and 70 deg/s. **E.** Same as *D*, but now for broad spiking V1 neurons in granular layers. **F.** Same as *D*, but now for broad spiking V1 neurons in infragranular layers. **G.** Same as *D*, but now for narrow spiking V1 neurons.

